# Spatially explicit capture–recapture models for relative density

**DOI:** 10.1101/2025.01.22.634401

**Authors:** Murray G. Efford

## Abstract

Spatially explicit capture–recapture (SECR) methods are used widely to estimate animal population density and related parameters. Maximum likelihood has been applied to two flavors of SECR model - a full model that includes absolute density as a spatially varying parameter, and a model conditional on the number of detected individuals that assumes uniform density. We show how the conditional model may be extended to estimate relative density as a function of habitat and other spatial variables. Unlike absolute density, coefficients of relative density are robust to bias from heterogeneous individual detection. The conditional model has the further advantage of seamlessly including individual covariates of detection, and it is faster to fit than the full model. The spatial absolute density surface may be derived from the conditional model fit. Population growth rate (relative density with respect to time) may also be estimated from a conditional model, given temporal data. These properties are demonstrated by examples and simulation. It is suggested that conditional-likelihood models, including those for relative density, play a central role in SECR.

## Introduction

Capture–recapture is used to estimate parameters of animal populations from detections of marked individuals. In spatially explicit capture–recapture (SECR) the population is conceived as a spatial distribution of individual activity centers (AC) (Borchers & Efford, 2008; Efford & Fewster, 2013; Royle et al., 2014; Borchers & Fewster, 2016; Efford, 2025a). The method is well-suited to data from automatic cameras, given that individuals can be distinguished by their natural marks, or from passive DNA samplers such as hair snares. By modeling the distance-dependent spatial detection process, SECR avoids the assumption of constant detection probability or effective sampling area that limits inference from raw counts used as population indices (Kéry & Royle, 2021 Chapter 1).

We are concerned with variation in population density relative to factors of biological interest. In the SECR model these factors are spatial or temporal covariates in a model for the intensity of a point process for AC. A spatial covariate may be any known attribute of points in the potential habitat; candidates include vegetation type, disturbance history, distance to human habitation, and the spatial coordinates themselves. Spatial variation in population density is of general interest, as shown by the frequency with which it appears in published SECR models (Tourani, 2022), and recent papers have identified specific reasons for modeling density. For example, correlated spatial variation in density and detection parameters causes bias in estimates of average density if not modeled (McLellan et al., 2023; Efford, 2025a). Variation in animal density provides important data for population management (e.g., Boulanger, Nielsen & Stenhouse, 2018; Lamb et al., 2018; Ferrão da Costa et al., 2025). As an aside, many papers include illustrations of so-called ‘density surfaces’ that are not based on modeled density, but rather aggregate the estimated sampling distributions of the AC of detected individuals; these have inferior statistical properties and are no substitute for a density model (Durbach et al., 2024).

The rich possibilities in SECR for modeling density across space and time have appeared to run up against constraints intrinsic to capture–recapture. For example, both spatial estimates of absolute density and non-spatial estimates of population size are negatively biased by unmodelled variation of detection parameters among individuals (e.g., Otis et al. 1978; Royle et al. 2013; Efford 2014; Efford and Mowat 2014; Efford 2025c). A workaround is to include in the model individual characteristics (size, age, sex, etc.) that may account for the variation. Individual covariates are readily included in models that use a likelihood conditional on the number of detected individuals. Such models in turn provide individual-specific estimates of capture probability or effective sampling area, leading by Horvitz–Thompson methods to robust estimates of non-spatial population size (Huggins 1989; Alho 1990) or population density (Borchers & Efford, 2008).

However, individual covariates have been considered incompatible with modeling spatial variation in SECR because that requires maximization of the full (unconditional) likelihood (Borchers & Efford, 2008; Borchers & Fewster, 2016; Sutherland, Royle & Linden, 2019), or an equivalent Bayesian method (Royle et al., 2014). This report shows that relative density (i.e. variation with respect to covariates) can be described by fitting the conditional model alone, leaving absolute density as an optional derived parameter, and that this approach has advantages for ecologists. Spatial pattern is usually represented in SECR models by a log-linear model for density as a function of location-specific covariates. For the absolute density *D* (**x**) at a point **x** (coordinates *x, y*) this takes the form

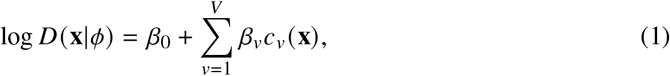

where the *c*_*v*_ (**x**) are *V* spatial covariates measured at **x** and the *β*s are coefficients to be estimated that together comprise the parameter vector *ϕ*. Categorical attributes are coded using indicator variables. The bases of regression splines may be used for fitting smooth trend surfaces (Borchers & Kidney, 2014). The logarithmic transformation ensures that the modeled density is always positive. The transformation is analogous to the ‘link’ function in generalized linear models (McCullagh & Nelder, 1989) and we use that term for convenience.

Relative density differs from absolute density by a constant factor that is given by the intercept *β*_0_. The benefits of focusing on relative density are demonstrated in this report by simulation and the analysis of example datasets. They may be summarized as follows. Fitting the conditional model is significantly faster than fitting the full model. While estimates of absolute density are biased by heterogeneity of detection parameters among individuals, heterogeneity has little effect on the estimated coefficients of relative density. Location-specific absolute density may be estimated as a derived parameter. Alongside any density covariates, the conditional model may include individual covariates of detection that improve model fit and potentially reduce bias in derived estimates of absolute density. Conditional models with spatial variation in density and models with individual covariates may be compared directly by information-theoretic criteria such as AIC as they are no longer in separate domains.

Change in population size or density over time (population growth rate) is central to demographic studies in ecology, and this is another application for conditional capture–recapture models. In non-spatial open population capture–recapture models, the emphasis has been on estimating per capita survival by the Cormack–Jolly–Seber (CJS) method that conditions on the first release of each marked animal (Lebreton et al. 1992). Conditioning on the total number of detected individuals in an open population model leads to estimates of both survival and recruitment (Pradel, 1996; Link & Barker, 2005; Schofield & Barker, 2016), where recruitment can be parameterized as the finite population growth rate *λ*. Efford & Schofield (2020) described a spatial version of the Pradel–Link–Barker (PLB) method in which *λ* measures change in population density i.e. *λ* is the *relative* density at consecutive discrete time steps. Temporal variation in density may also be inferred from a chain of samples (‘sessions’) analysed without regard to individual survival, as an extension of closed-population SECR (Efford, Borchers & Byrom, 2009; Borchers & Fewster, 2016; Sutherland, Royle & Linden, 2019; Efford, 2025a). A conditional multi-session closed model is proposed here that leads directly to estimates of the session-specific *λ*.

Concluding remarks emphasize the central place of relative-density methods in SECR. Many questions can be answered directly by fitting the relative density model, and absolute density is available as a derived parameter if needed. Certain limitations of the method are acknowledged.

## Conditional likelihood for relative density

The full SECR likelihood has the form

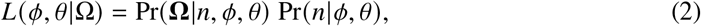

where Ω is the set of *n* observed spatial detection histories, *ϕ* is a vector of density parameters and *θ* is a vector of detection parameters (Borchers & Efford, 2008). The first factor on the right of Equation (<u>2</u>) concerns the joint probability of the *n* observed detection histories. The second factor concerns the number of detected animals *n*. When individuals are detected independently of each other, the first factor may be expanded to

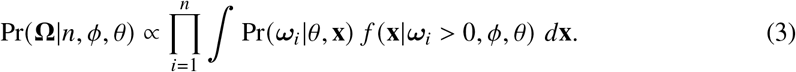

The actual location of each AC is unknown, so the location **x** is ‘marginalized’ out of the likelihood by integrating over all possible locations in two dimensions. A critical factor is the probability density *f* (**x**|***ω***_*i*_ *>* 0, *ϕ, θ*) for the AC location **x** of individual *i*, given that the individual was detected (*ω*_*i*_ *>* 0). That probability density has components for (i) the spatial distribution of the population, modeled with the density function *D* (**x**|*ϕ*) from Equation (1), and (ii) the probability *p*_·_ (**x**|*θ*) that an individual with AC at **x** appears in the sample. Formally:

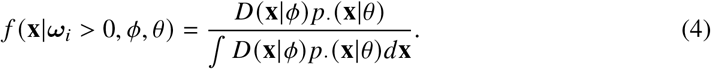

The probability *p*_·_ (**x**|*θ*) combines the probabilities of detection at each detector, each possibly compounding detection over multiple occasions (Borchers & Efford, 2008; Efford, Borchers, & Byrom, 2009; Appendix S1). If density is uniform then *D* (**x**|*ϕ*) cancels out of Equation (4) and maximizing the likelihood conditional on *n* in Equation (3) provides an estimate of *θ* (Borchers & Efford, 2008). However, if density is not uniform then *D* (**x**|*ϕ*) does not cancel out of Equation (4) and the spatial variation in density must be included to obtain an unbiased estimate of *θ*. This is not as onerous as it sounds.

We define relative density by *D*^′^(**x**|*ϕ*^−^) ≡ *k*^−1^*D* (**x**|*ϕ*), where *ϕ*^−^ is a vector with one fewer coefficients than *ϕ* and *k*^−1^ is a constant of proportionality. For the log-linear model, *k* = exp(*β*_0_) and *ϕ*^−^ is simply *ϕ* without the intercept. We simplify and maximize the conditional likelihood in Equation (3) to estimate both the detection parameters *θ* and the coefficients of relative density *ϕ*^−^. Cancelling the constant *k* yields the conditional likelihood

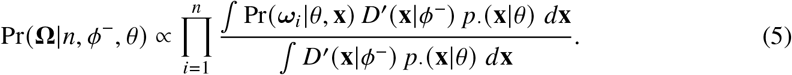

### Individual covariates

Spatial detection is governed by the parameter vector *θ* that typically comprises the parameters *λ*_0_ and *σ* for the intercept and spatial scale of a function relating detection hazard to distance (Appendix S1). This use of *λ*_0_ is unrelated to the conventional use of *λ* for population growth rate. Known predictors of variation, such as weather or trap type, are included by specifying a linear submodel for either parameter on its link scale, as for density in Equation (1). Each submodel may also include predictors that vary among individuals (‘individual covariates’), but these raise issues as we describe next.

To include individual covariates for detection parameters in the full likelihood it is necessary to model their distribution in the population. That is likely to differ from the distribution in the sample due to sampling bias. The shape of the covariate distribution must be specified and each evaluation of the likelihood requires integration over potential values of the covariate (e.g., Pollock, 2002). This is feasible for a single binary variable (e.g., sex) as the only parameter is the mixing proportion (e.g., sex ratio). For multi-level or continuous individual covariates the added model complexity and computational burden have discouraged implementations using the full likelihood. The problem is bypassed in Bayesian SECR using the complete data likelihood, as the population distribution of each individual covariate is included in the model (Link & Barker, 2010); an arbitrary shape is specified for the distribution (e.g., normal) and integration happens implicitly.

The problem is also bypassed by maximizing the conditional likelihood, without the need to specify the population distributions of covariates (Huggins, 1989; Alho, 1990). The conditional model concerns only detected individuals whose covariates are observed directly. Thus maximizing the conditional likelihood allows us simultaneously to fit models for both relative density and covariate effects on individual detection. Further, Horvitz-Thompson-like methods for inferring absolute population size or density from a sample adjust for sampling bias with respect to covariates.

### Absolute density as a derived parameter

Absolute density may be computed directly from a fitted model for relative density. Derived estimates of density are the same as those from maximization of the full likelihood if the AC follow a Poisson distribution in two dimensions, whether that is homogeneous or inhomogeneous (see Discussion).

Given uniform density, the effective sampling area *a* is a function of *θ* that may include a vector of covariates **z**_*i*_ for each individual *i*, hence *a* (*θ*, **z**_*i*_) = ∫*p*_·_ (**x**|*θ*, **z**_*i*_) *d***x**. This leads to the Horvitz-Thompson-like estimate of the uniform density 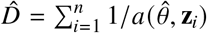 (Borchers & Efford, 2008). For non-uniform density, observe that when *p*_·_ (**x**|*θ*) does not depend on individual covariates, E(*n*) = *k* ∫*D*^′^(**x**|*ϕ*^−^) *p*_·_ (**x**|*θ*) *d***x**, and 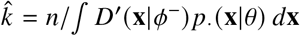, where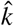 provides the missing intercept of the absolute density model (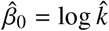). When detection parameters depend on individual covariates the factor *n* is replaced by summation over *n*:

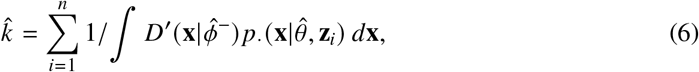

The sampling variance of *β*_0_ and its covariance with other *β*s may be estimated by the delta method using numerical estimates of the gradient with respect to the estimated 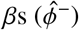 For uniform density *D*^′^(**x**|*ϕ*^−^) takes the value 1, and ∫ *D*^′^(**x**|*ϕ*^−^) *p*_·_ (**x**|*θ*, **z**_*i*_) *d***x** collapses to the effective sampling area of individual *i*. Equation (6) is then the previous Horvitz-Thompson-like estimate. Further, a derived estimate of the absolute density at a point **x** is

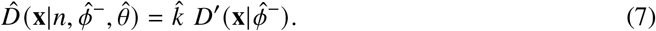

### Multi-session conditioning

When the data span multiple independent sub-populations (‘sessions’), and the density model describes variation among them such as a temporal trend, conditioning is on the total number observed, as in PLB models (Efford & Schofield, 2020). The conditional likelihood is then the product of the *T* session-specific likelihoods and a factor for the session-specific counts *n*_*t*_, modelled as multinomial with size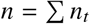:

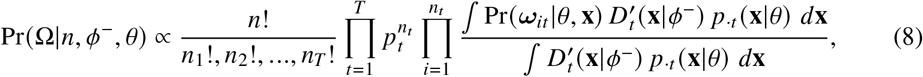

where subscript *t* indicates the session. The multinomial probabilities *p*_*t*_ are estimated from the normalized expected number of detections in each session, using session-specific derived estimates of absolute density.

In order to parameterize the temporal model as population growth rate we model the cumulative change in density:

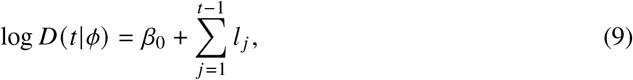

where *β*_0_ is the initial log density and *l*_*t*_ is a linear predictor such that *λ*_*t*_ = exp *l*_*t*_. In the simplest model, with equal time intervals and session-specific *λ*_*t*_, each *l*_*t*_ corresponds to one *β* and the parameter vector for the conditional model is *ϕ*^−^ = (*β*_1_, *β*_2_, …, *β*_*T*−1_). Otherwise, there may be a linear submodel for the *l*_*t*_ in terms of a reduced set of *β*s, and the *l*_*t*_ may be scaled by time interval.

## Simulations

Simulations were used to evaluate the performance of conditional likelihood models when the detection parameter *λ*_0_ varied among individuals. Spatial pattern was generated as a Gaussian random field whose intensity *c*_1_(**x**) was scaled to (0,1) and treated as a known covariate. Expected density was modeled by log *D* (**x**) = *β*_0_ + *β*_1_*c*_1_(**x**), where *β*_0_ = log(0.2) and *β*_1_ = log(10); density therefore ranged between 0.2*σ*^−2^ and 2*σ*^−2^ depending on the covariate value at **x**. AC were generated from an inhomogeneous Poisson distribution with the modeled density, and detections with a hazard-halfnormal function were simulated on a 10 × 10 square grid of binary proximity detectors at 2-*σ* spacing (Fig. 1). A conditional SECR model was fitted with the same structure as the generating model. The fitted model estimated the coefficient 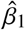directly, and 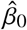 was derived using Equation (6). The intercept *λ*_0_ of the exponential hazard function was allowed to vary among individuals according to a log-normal distribution with mean 0.2 and CV 0, 0.2, 0.4, 0.6 or 0.8. Simulations were repeated 2000 times for each level of CV(*λ*_0_). The average number of individuals detected declined with increasing CV from *n* = 265 individuals (SD = 71.6) (CV(*λ*_0_) = 0) to 232 (SD 60.7) (CV(*λ*_0_) = 0.8), while the average number of detections per individual increased from 1.77 to 1.97. Results were expressed as the average absolute bias of each coefficient (absolute bias on the log scale approximates relative bias on the natural scale). R code and detailed results are given in section ‘MhS’ of Efford (2025c).

**Figure 1.**
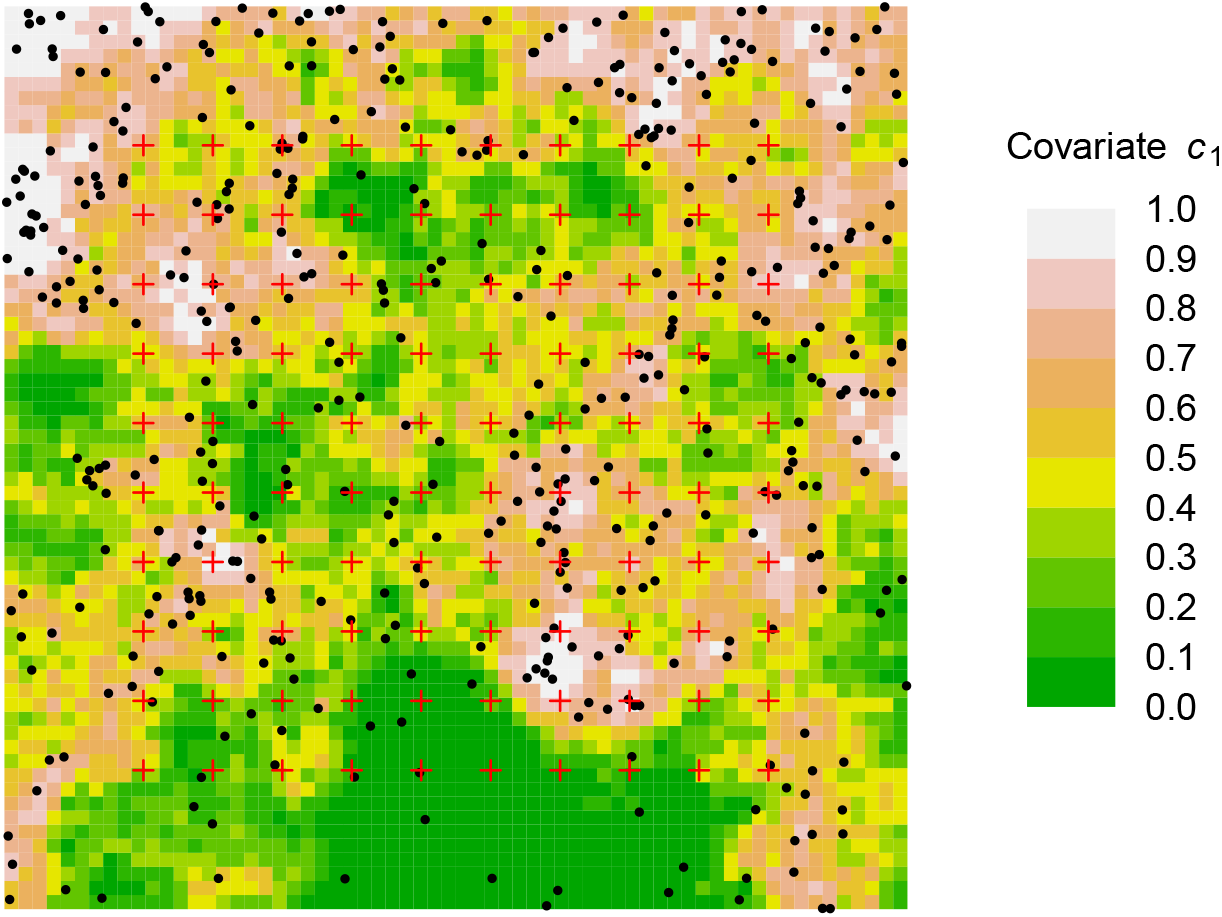
Example of simulated spatial covariate *c*_1_ and activity centres (black dots) generated from inhomogeneous Poisson model for intensity as a log-linear function of the covariate. Detector sites are shown as red crosses.

Estimates of 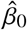, the intercept that determines absolute density, were strongly biased by unmodelled variation in detection, but, estimates of relative density (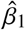) were unaffected (Fig. 2). Execution time for the conditional likelihood fit (3 parameters) averaged 59% to 70% of that for the full likelihood (4 parameters). The absolute difference between estimates of *β*_1_ from the full and conditional models was less than 0.0003 in all simulations.

**Figure 2.**
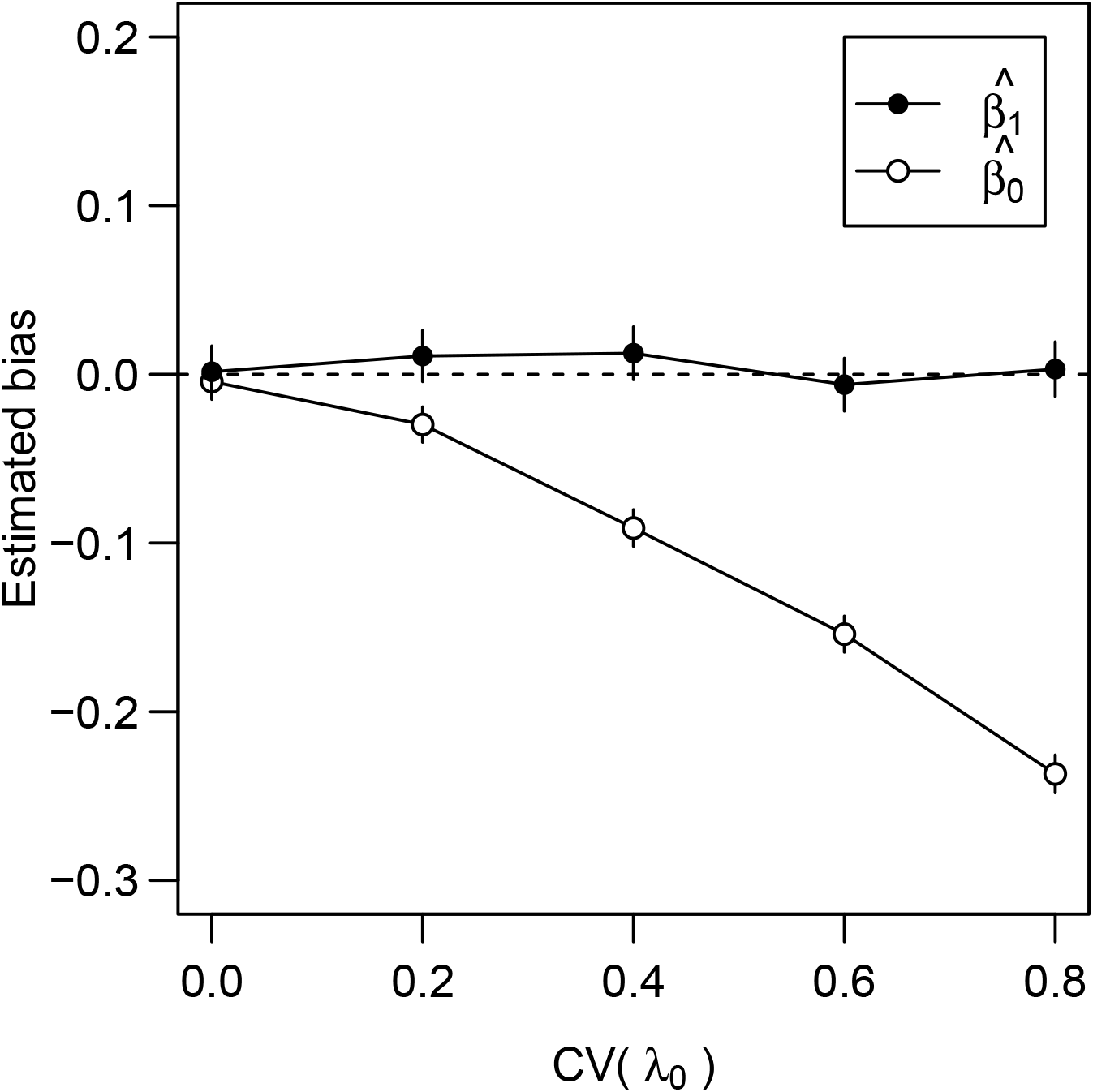
Bias of estimated coefficients of SECR model for density in relation to unmodeled individual heterogeneity in baseline detection parameter *λ*_0_. The log-linear spatial density model had intercept *β*_0_ and slope *β*_1_ with respect to covariate *c*_1_ (Fig. 1). Simulated data: average of 2000 replicates with 95% CI.

Simulations were also conducted to assess the effect of variation in *λ*_0_ on estimates of density and population growth rate from temporal data. The pattern of bias followed that for spatial pattern (density biased, trend unbiased) so detailed results are not presented here; they are available in section ‘MhT’ of Efford (2025c).

## Example. Spatial distribution of skinks

Data were analysed from a study of diurnal lizards in the Upper Buller Valley, South Island, New Zealand. Pitfall traps (covered sunken cans baited with a morsel of fruit in sugar syrup) were set in two large grids, each with 231 traps about 5 m apart and operated intermittently through 1995–1996. Traps were checked daily and individuals were marked uniquely by toe clipping. We selected data on one species (*Oligosoma infrapunctatum*) trapped for 3 days in November 1995. Ground cover and vegetation were recorded in a 1-m radius plot at each trap site. Each site was assigned to one of two habitat classes: ‘low cover’ (open with bare ground or low-canopy vegetation and grasses; 52.6% of sites) and ‘high cover’ (more-closed vegetation with a higher canopy; 47.5% of sites). A subset of the data was used by Efford & Fewster (2013) and the full dataset is included in Efford (2025b), where it is described further.

SECR models were fitted conditional on the number of trapped individuals, *n* = 253. Individuals varied greatly in body size measured as snout-vent length (SVL). This was incorporated as a predictor of the detection parameters *λ*_0_ and *σ* either as a log-linear effect or as non-linear spline smooths with 3 or 5 degrees of freedom. Each candidate detection model was fitted with and without an effect of habitat class on local density. The model with lowest AIC included both the habitat effect on density and a non-linear effect of body size on *λ*_0_ (Fig. 3). A conditional-likelihood habitat model without individual covariates model fit in 58% of the time taken by a matching full-likelihood model. Details are in Appendix S2.

**Figure 3.**
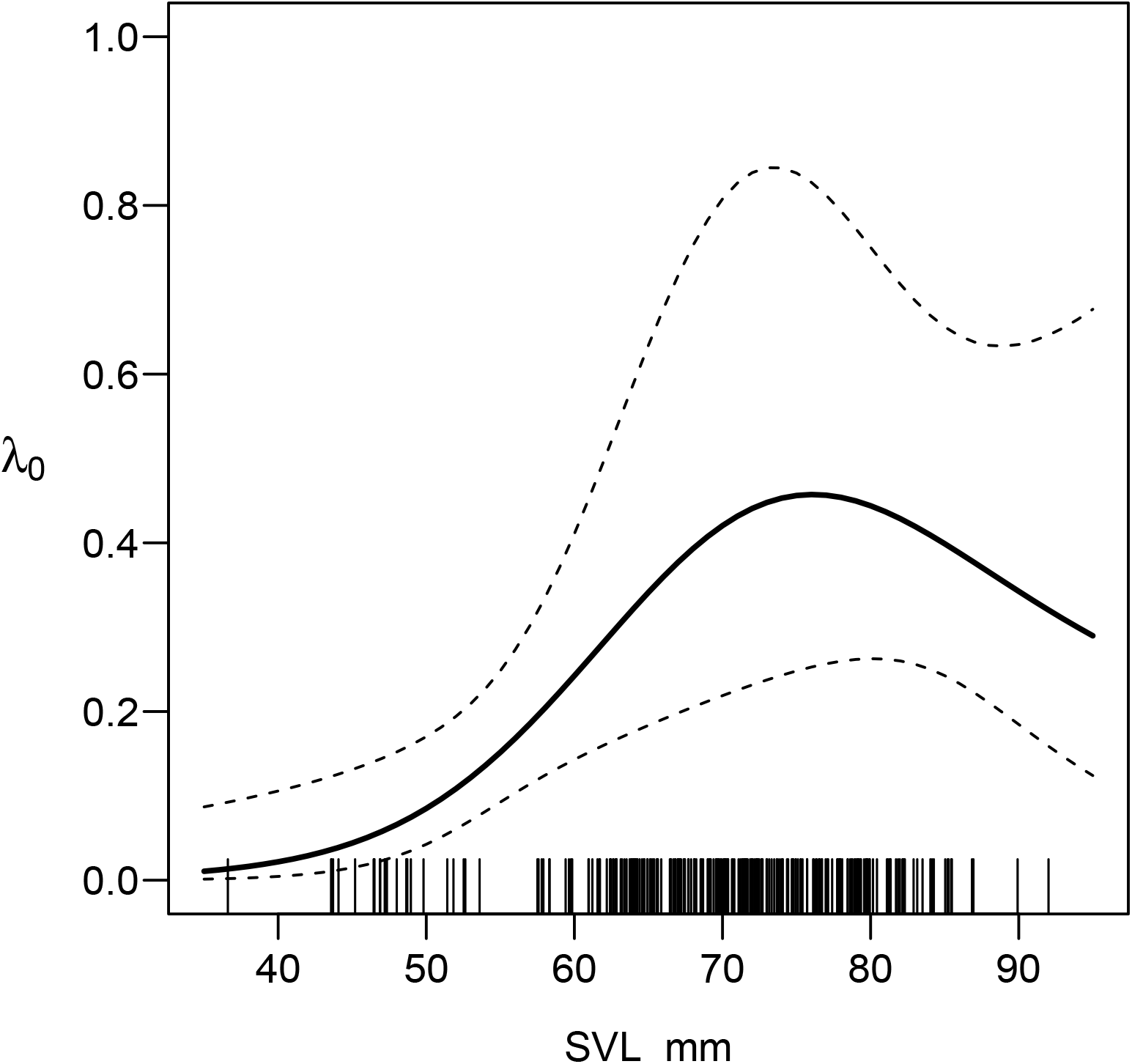
Fitted relationship between the baseline detection hazard *λ*_0_ and the individual covariate snout-vent length (SVL) of pitfall-trapped skinks *Oligisoma infrapunctatum*. Estimates from regression splines fitted in a conditional-likelihood SECR model. Rug plot indicates distribution of observed SVL.

*Oligosoma infrapunctatum* was concentrated at sites with higher vegetation: density in the ‘high’ class exceeded density in the ‘low’ class by a factor of exp *β*_1_ = 43.2, regardless of whether the model allowed for the effect of body size on detection. Allowing for body size considerably increased the estimate of density, but the precision of density estimates was much reduced (Table 1).

**Table 1.**
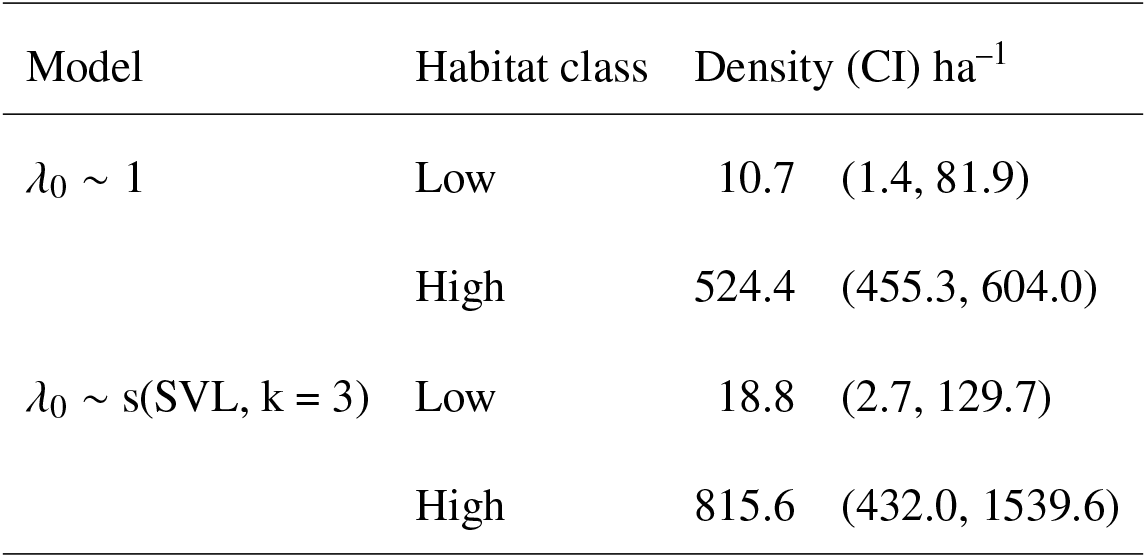
Estimates of habitat-specific density of the skink *Oligosoma infrapunctatum* from model with no individual covariate of detection hazard *λ*_0_, and a model with spline smooth of snout-vent length *λ*_0_ ∼ s(SVL, k = 3).

## Example. Brushtail possum population dynamics

The conditional-likelihood method for relative density through time (i.e. population growth rate *λ*) is illustrated with data from a multi-year trapping study of the brushtail possum *Trichosurus vulpecula* in New Zealand. A population in mixed forest was monitored by live trapping (Efford & Cowan, 2004). Analysis focused on June data from 1996 to 2005, a period of constant trapping effort (167 traps were set at the same sites at 30-m spacing for 5 nights). In total, 698 individuals were captured 4408 times. Annual population growth rate was estimated by multi-session closed-population SECR, parameterized as in Equation 9 of the main text. Direct estimates of *λ* from the conditional model parameterized as relative density were indistinguishable from those computed as the ratio of successive density estimates from maximization of the full-likelihood (Fig. 4). The year-specific conditional-likelihood model fit in 78% of the time taken by a matching full-likelihood model. The direct approach enables further modeling of *λ*. Models were compared with fixed *λ*_*t*_ = 0 (constant density), constant *λ*_*t*_, session-specific *λ*_*t*_, and smooths with varying degrees of freedom. Appendix S3 gives further details and results, including comparisons with open-population (PLB) models fitted to the same data.

**Figure 4.**
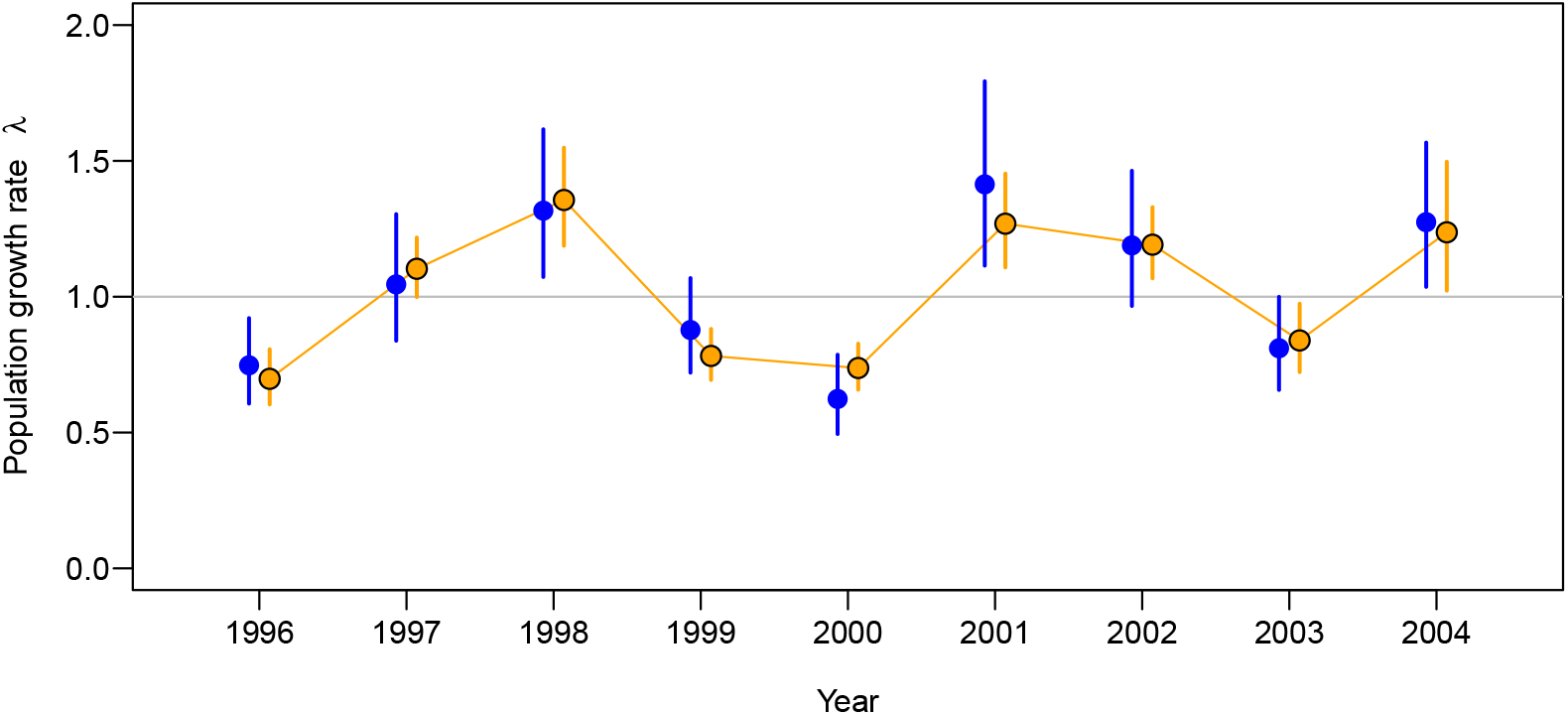
Annual population growth rate *λ* of brushtail possums *Trichosurus vulpecula*. The model for *λ* used either a 9-level factor for year-specific estimates (blue) or a spline smooth with *k* = 7 (orange, joined). Bars are 95% confidence intervals. The smooth model had lower AIC and shorter intervals, similar to those from an open population model (Appendix S3). Estimates of *λ* as the ratio of successive session-specific estimates of density from a full-likelihood model exactly overlaid the session-specific conditional-likelihood 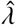 and are not shown.

### Implementation

The number of identifiable coefficients for relative density in the conditional model (*ϕ*^−^) is one less than the number for absolute density in the full model (*ϕ*). Given software for the full likelihood, the conditional likelihood is maximized by omitting the likelihood component for *n* and fixing the intercept *β*_0_ to zero, assuming a log link for density. One may also use an identity link truncated at zero (Efford & Fewster, 2013); then *k* = *β*_0_ and *ϕ*^−^ is *ϕ* with the intercept removed and remaining coefficients divided by *β*_0_. For the log link, estimated coefficients of the Poisson relative density model are identical within numerical error to the corresponding coefficients of the full model. For the identity link, the estimated coefficients equal those from the full model divided by its intercept. The density intercept on the natural scale may be derived from the density-weighted effective sampling areas of Equation (6).

Estimation of relative density by maximizing the conditional likelihood is included in recent versions of the R package ‘secr’ (R Core Team, 2025; Efford, 2025b). Integration is performed in ‘secr’ by summing the integrand over a set of pixels for which its value is non-negligible, known as the habitat mask. The conditional likelihood in Equation (5) is maximized numerically whenever the CL argument of function ‘secr.fit’ is TRUE and a model formula is specified for density. The usual asymptotic variance-covariance matrix for coefficients in the model (*ϕ*^−^, *θ*) is obtained from the negative inverse Hessian evaluated at the MLE. Functions ‘derivedDcoef’ and ‘derivedDsurface’ implement Equations (6) and (7) respectively. The parameterization in Equation (9) is selected with the ‘Dlambda’ details option.

## Discussion

A model for the distribution of animal activity centers may be split into a model for the density relative to spatial or temporal covariates and a scalar that converts relative density to absolute density. Relative density captures effects of ecological interest and is estimated efficiently by maximizing the likelihood conditional on *n*, the number of observed individuals. This report emphasises the central role and advantages of the conditional model, while acknowledging some limitations.

Spatial variation in relative density is fully captured by a conditional model. Coefficients for both detection (*θ*) and covariate effects on density (*ϕ*^−^) do not differ between equivalent full- and conditional-likelihood models when the AC come from an inhomogeneous Poisson process, as in the simulations. The key here is that the statistic *n* is then Poisson (e.g., Efford & Fletcher, 2025), and hence *n* is S-ancillary for the model coefficients in *θ* and *ϕ* (Schofield & Barker, 2016).

S-ancillarity means that *n* need not be modeled, and that derived estimates of absolute density from the conditional model are identical to full likelihood estimates, within the limits of the maximization algorithm. This property does not hold exactly when the number of AC in a specified region is assumed to be fixed rather than Poisson; then the fitted model for *n* is binomial, and *n* is no longer S-ancillary for the coefficients (Schofield & Barker, 2016).

Simulations also demonstrated that the coefficients of *relative* density (i.e. *ϕ*^−^, excluding *β*_0_) are robust to unmodelled individual heterogeneity. This is true whether they are estimated by maximizing the full or conditional likelihoods. However, estimates of *β*_0_ and hence *absolute* density from either model are biased by such heterogeneity. If absolute density is required, a partial solution unique to the conditional model is to include individual covariates in submodels for the detection parameters. This was demonstrated in the skink example, although confidence intervals of the resulting estimates were wide.

Ecological investigations using SECR typically compare multiple models by information-theoretic criteria such as AIC to find a ‘best’ model or to provide weights for model averaging. Comparison is possible only within a class of model (full or conditional). The greater scope of conditional models, now including both relative density and individual covariates, suggests that they should be used for all such comparisons. Estimating one fewer parameters than in the full model leads to faster model fitting, with time savings of 42% and 22% in the examples.

The effectiveness of the conditional model for estimating spatial variation in relative density has a direct equivalent in the Pradel–Link–Barker (PLB) open population model that also conditions on *n* (Schofield & Barker, 2016; Efford & Schofield, 2020). Temporal variation of the finite population growth rate in PLB models is analogous to spatial variation in relative density. A temporal series of closed-population samples may be analysed jointly by SECR, and it is convenient to parameterize the joint model in terms of population growth rate, as in the brushtail possum example (Fig. 4). Marescot et al. (2011) drew attention to the robustness and utility of non-spatial estimates of population growth rate. Simulations reported separately (Efford, 2025c) confirmed the robustness to individual heterogeneity of the population growth rate estimated from a temporal series of closed-population samples. This mirrors the robustness of relative density with respect to spatial covariates (Fig. 2).

### Limitations

A case has been made for using SECR to model spatial variation in density, but there are limits to the ability of SECR of whatever flavor to resolve patterns and attribute variation to habitat covariates. The underlying Poisson point process model is stochastic and large samples are required to resolve patterns. The spatial detection process (scale *σ*) blurs the picture, especially when the study area is small relative to *σ*. The precision of habitat-specific estimates of absolute density can be poor, and is exacerbated by uncertainty in the detection model, as in the skink example.

Derived estimates of absolute density are readily computed by the Horvitz-Thompson method from any CL model. However, their variances must be obtained by the delta method which relies on a numerical estimate of the gradient vector for density with respect to the fitted coefficients (Huggins, 1989). Profile likelihood confidence intervals are not an option.

It is not possible to treat absolute density as a derived parameter when another component of the likelihood depends upon it. Specifically, the conditional model does not allow the spatial scale of detection *σ* to be a function of absolute density (Efford et al., 2016).

Variance estimates for population growth rate from a sequence of closed populations assume that each comprises an independent Poisson sample of AC. Survival of some individuals, each keeping its AC, violates this assumption and seems likely to cause a spurious improvement in precision, relative to an open population model that includes survival. However, the reverse appeared to be the case in the brushtail possum example (Appendix S3), possibly because survival reduced the component to be estimated. Further investigation is warranted.

## Conclusion

Pollock (2002) and Royle (2009) expressed unease about the use of the conditional likelihood in closed-population capture–recapture on the abstract ground that the parameter of interest (population size in their case) was not in the model. However, there are few negative consequences in practice (preceding section). The conditional SECR model allows relative density to be modeled with respect to covariates despite the omission of absolute density. Fitting is fast and models with any combination of spatial and individual covariates may be compared by AIC. Attention is focused on the robustly estimated coefficients of the density model rather than the non-robust intercept that determines absolute density. Models fit by maximizing the conditional likelihood are sufficient to address questions regarding spatial or temporal variation in density, and they are a logical first step when the goal is to estimate absolute density.

## Supporting information

Appendix S1 Probability of detection

Appendix S2 Skink example

Appendix S3 Brushtail possum example

## Acknowledgments

I thank the many people who helped collect field data. The Lake Station skink study was a collaboration with Bruce Thomas, Nick Spencer, Rob Mason and Peter Williams. Contributors to the possum study are listed in Appendix S3. I acknowledge the use of New Zealand eScience Infrastructure (NeSI) high performance computing facilities URL https://www.nesi.org.nz. I thank Matt Schofield, Pierre Dupont and two anonymous referees for their comments on drafts.

