## Appendix S1 Probability of detection for "Spatially explicit capture–recapture models for relative density"

### Appendix S1. Model for overall probability of detection

For independent events,

$$p_{\cdot}(\mathbf{x}|\theta) = 1 - \prod_{k}^{k=K} \prod_{j}^{j=J} [1 - p_{jk}(\mathbf{x}|\theta)], \tag{1}$$

where $p_{jk}(\mathbf{x}|\theta)$ is the probability of detection on occasion $j$ at detector $k$ given the vector of parameters $\theta$. For present purposes we assume the $p_{jk}$ are all governed by a distance-detection function such as the halfnormal $p_k(\mathbf{x}|\theta) = g_0 \exp[-d_k^2/(2\sigma^2)]$ where $d_k$ is the distance between AC location $\mathbf{x}$ and detector $k$, and parameter vector $\theta = (g_0, \sigma)$. Then

$$p_{\cdot}(\mathbf{x}|\theta) = 1 - \prod_{k}^{k=K} [1 - p_k(\mathbf{x}|\theta)]^J. \tag{2}$$

Other distance-detection functions are the negative exponential and hazard rate (Efford 2025). Each function has an equivalent in which the hazard of detection $\lambda(d)$ is modeled instead of the probability of detection (hence 'hazard halfnormal', 'hazard hazard-rate', and 'hazard negative exponential'). The probability of detection from a hazard model is $p_k(\mathbf{x}|\theta) = 1 - \exp -\lambda(d)$.
