## Appendix S2 Skink example for "Spatially explicit capture–recapture models for relative density"

### Appendix S2. Spatial distribution of skinks

Skinks (*Oligosoma infrapunctatum*) were captured in pitfall traps checked daily for 3 days. Individuals were marked uniquely by clipping one toe on each foot. The capture data are provided as object infraCH in the R package 'secr' (Efford 2025a); only the second session was used (14–17 November 1995). Pitfall traps are multi-catch traps, i.e. multiple skinks could be caught at one time, but each skink could only appear in one trap per day (infrequent repeat captures were discarded). The activity centers (AC) of captured skinks were assumed to lie within 15 m of a trap: the habitat mask was constructed using a 15-m buffer around the traps and discretized as 2-m pixels. Each pixel was assigned to a 'low cover' or 'high cover' habitat class as described in the main text.

Analyses were run in R 4.5.1 on a x86_64 machine in Windows 11 using package secr version 5.3.0 and 18 cores. Data, R code, and fitted models are on Zenodo (Efford 2025b).

Several distance-detection functions were compared in exploratory analyses (i.e. with all other parameters held constant).

**Table S 1:** Evaluation of candidate functions for detection hazard as function of distance

| Function | npar | logLik | ΔAIC | AICwt |
|---|---|---|---|---|
| HHR hazard rate | 3 | –1340.571 | 0.000 | 0.9252 |
| HVP variable power | 3 | –1343.086 | 5.030 | 0.0748 |
| HEX exponential | 2 | –1350.102 | 17.062 | 0.0000 |
| HHN halfnormal | 2 | –1378.224 | 73.306 | 0.0000 |

The 3-parameter hazard-rate function had the lowest AIC (Table S1) and was used in later model comparisons of more complex models. This detection function has the form

$$\lambda(d) = \lambda_0[1 - \exp\{-(^d/_\sigma)^{-z}\}], \quad (1)$$

where $\lambda(d)$ is the hazard of detection for an activity center (AC) at distance $d$ from a trap and $\lambda_0, \sigma$ are the intercept and spatial scale as in the more common half-normal model, and $z$ is a shape parameter (assumed constant in all analyses).

The main analyses compared models with and without an effect of habitat class for various linear or smooth relationships between body size (snout-vent length SVL) and the detection parameters $\lambda_0$ and $\sigma$ (Table S2). SVL was binned in 5-mm intervals to reduce computation time (hence 'SVL5'). The spline smooth s() follows R package 'mgcv' (Wood, 2017); s(SVL5, k = 3) indicates a spline smooth of SVL5 with k–1 = 2 degrees of freedom.

**Table S 2:** Candidate skink models.

| Code | $D$ | $\sigma$ | $\lambda_0$ |
|---|---|---|---|
| hhr | D~1 | sigma~1 | lambda0~1 |
| svl | D~1 | sigma~SVL5 | lambda0~SVL5 |
| svl.s3 | D~1 | sigma~s(SVL5, k = 3) | lambda0~s(SVL5, k = 3) |
| svl.s5 | D~1 | sigma~s(SVL5, k = 5) | lambda0~s(SVL5, k = 5) |
| svl.s3l | D~1 | sigma~1 | lambda0~s(SVL5, k = 3) |
| svl.s5l | D~1 | sigma~1 | lambda0~s(SVL5, k = 5) |
| hab.hhr | D~habclass | sigma~1 | lambda0~1 |
| hab.svl | D~habclass | sigma~SVL5 | lambda0~SVL5 |
| hab.svl.s3 | D~habclass | sigma~s(SVL5, k = 3) | lambda0~s(SVL5, k = 3) |
| hab.svl.s5 | D~habclass | sigma~s(SVL5, k = 5) | lambda0~s(SVL5, k = 5) |
| hab.svl.s3l | D~habclass | sigma~1 | lambda0~s(SVL5, k = 3) |
| hab.svl.s5l | D~habclass | sigma~1 | lambda0~s(SVL5, k = 5) |

**Table S 3:** Comparison of skink models

| | npar | logLik | ΔAIC | AICwt |
|---|---|---|---|---|
| hab.svl.s3l | 6 | -1240.94 | 0.00 | 0.69 |
| hab.svl.s5l | 8 | -1240.70 | 3.52 | 0.12 |
| hab.svl.s3 | 8 | -1240.72 | 3.56 | 0.12 |
| hab.svl.s5 | 12 | -1237.37 | 4.86 | 0.06 |
| hab.svl | 6 | -1244.67 | 7.46 | 0.02 |
| hab.hhr | 4 | -1261.29 | 36.70 | 0.00 |
| svl.s3l | 5 | -1320.01 | 156.14 | 0.00 |
| svl.s3 | 7 | -1319.44 | 159.00 | 0.00 |
| svl.s5l | 7 | -1319.79 | 159.70 | 0.00 |
| svl.s5 | 11 | -1316.62 | 161.37 | 0.00 |
| svl | 5 | -1324.06 | 164.25 | 0.00 |
| hhr | 3 | -1340.57 | 193.26 | 0.00 |

**Table S 4:** Estimates from AIC-best fitted model. $\lambda_0$ is predicted for the median SVL (70 mm).

| | estimate (SE) | | 95% CI |
|---|---|---|---|
| D Low ha$^{-1}$ | 18.9 | (24.1) | (2.7, 129.7) |
| D High ha$^{-1}$ | 815.6 | (271.5) | (432.0, 1539.6) |
| $\lambda_0$ | 0.371 | (0.099) | (0.222, 0.619) |
| $\sigma$ m | 3.50 | (0.42) | (2.77, 4.42) |
| $z$ | 3.60 | (0.25) | (3.14, 4.13) |

**Table S 5:** Relative bias in simulations of habitat-specific density of the skink *Oligosoma infrapunctatum* from two models. True density Low habitat 18.9 $ha^{-1}$, High habitat 815.6 $ha^{-1}$. Average of 100 replicates.

| Model | Habitat class | RB($\hat{D}$) (SE) |
|---|---|---|
| $\lambda_0 \sim 1$ | Low | –0.175 (0.059) |
| | High | –0.082 (0.005) |
| $\lambda_0 \sim$ s(SVL, k = 3) | Low | –0.073 (0.066) |
| | High | 0.017 (0.008) |

### Timing

Timing was compared between a full-likelihood fit of the habitat-class model using the 3-parameter hazard-rate detection function with no individual covariates (5 parameters), and a conditional-likelihood fit of the same model (4 parameters). The conditional fit took 8.13 seconds compared to 13.99 seconds for the full likelihood.

### Simulations

A small simulation study was performed to assess the effect of size-related heterogeneity in $\lambda_0$ on estimates of habitat-specific population density, given estimates from the AIC-best fitted model 'hab.svl.s3l'. Distributions of AC were generated across the skink habitat mask at the estimated densities. Individual values of $\lambda_0$ were drawn with replacement from observed SVL values with probabilities weighted inversely by the SVL-specific effective sampling area $a$. Inverse weighting corrects for underrepresentation of small skinks in the sample. Sampling with the two grids (462 traps) was simulated for 3 days using the HHR detection function and $\sigma = 3.5$ m and $z = 3.6$ as estimated. Two models were fitted to each dataset, hab.hhr and hab.svl.s3l in Table S2.

Results are expressed as the bias of each habitat-specific density relative to the true density for the habitat (Table S5). Estimates for the High habitat with the correct model were essentially unbiased, while the misspecified null model ($\lambda_0 \sim 1$) showed a –8% bias. Estimates for the Low habitat were imprecise, presumably because of the very low density, but showed a similar trend.
