## Appendix S3 Brushtail possum example for "Spatially explicit capture–recapture models for relative density"

### Appendix S3. Brushtail possum population dynamics

A population of brushtail possums *Trichosurus vulpecula* was monitored in a New Zealand forest (Efford & Cowan 2004). Possums were trapped in cage traps and marked with ear tags and numbered tattoos. Analysis focused on June data from 1996 to 2005, a period of constant trapping effort (167 traps were set at the same sites at 30-m spacing for 5 nights). In total, 698 individuals were captured 4408 times. Data, R code, and fitted models are on Zenodo (Efford, 2025).

Annual population growth rate was estimated by conditional closed-population SECR, parameterized as in Equation 9 of the main text, and by the equivalent open-population spatial method (Efford & Schofield, 2020). Estimates were very similar, but confidence intervals were much shorter for the open-population estimates (Fig. S1). Annual survival was high (0.56–0.86) and many individuals from year $t$ were known to be alive in year $t+1$, reducing uncertainty in the open-population estimates. For interest, we contrived an open-population analysis that discarded information on survival by assigning a year-specific identity to each individual and fixing the survival rate at zero; then the PLB estimates were numerically indistinguishable from the closed-population estimates.

The possum population showed no net change over 1996–2005 (fixed-$D$ same likelihood as constant-trend model Table S1).

Confidence intervals for $\hat{\lambda}$ were shorter using the conditional open-population (PLB) approach than with the year-specific closed-population model, presumably because many individuals survived between sessions (Fig. S1). However, the difference largely disappeared, with the AIC-best closed population model, which smoothed $\hat{\lambda}$.

Analyses were run in R 4.3.2 on a x86_64-pc-linux-gnu platform in Rocky Linux 9.4 using packages secr version 5.3.0, openCR version 2.2.7, and 60 cores. Computation was faster for the comparable closed models. The timings for a fully year-specific model were: closed population full likelihood 326 s (12 parameters), closed population conditional likelihood 254 s (11 parameters), open population spatial PLB 2046 s (20 parameters).

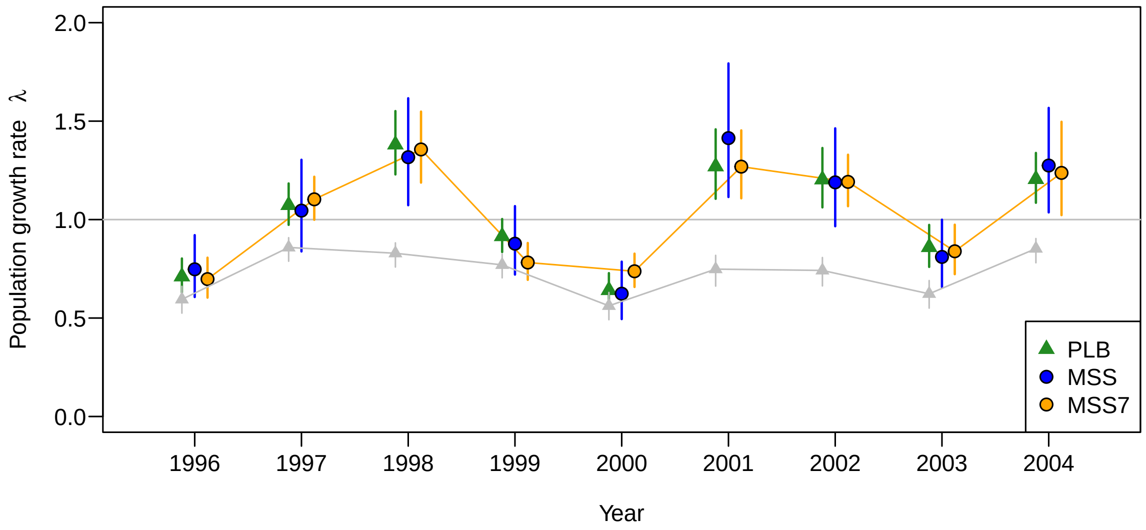


**Fig. S 1:** Annual population growth rate of brushtail possums *Trichosurus vulpecula* over 1996–2005 in New Zealand mixed forest. Estimates from spatial closed-population analysis, either fully year-specific (MSS) or with smoother (MSS7), compared to spatial open-population capture–recapture (PLB). Bars show 95% CI. Grey points indicate the survival component of the PLB estimates.

**Table S 1:** Comparison of conditional likelihood models for population growth rate of brushtail possums 1996–2005. The smooth year model with 7 knots had the lowest AIC and is plotted in the main text.

| $\lambda$ model | npar | logLik | dAIC | AICwt |
|---|---|---|---|---|
| s(year, k = 7) | 8 | -15915.67 | 0.00 | 0.82 |
| year-specific | 11 | -15914.18 | 3.01 | 0.18 |
| s(year, k = 3) | 4 | -15934.82 | 30.28 | 0.00 |
| s(year, k = 5) | 6 | -15932.84 | 30.32 | 0.00 |
| fixed $\lambda$ = 1.0 | 2 | -15938.23 | 33.11 | 0.00 |
| constant $\lambda$ | 3 | -15938.21 | 35.06 | 0.00 |

### Acknowledgments

Many people helped with the study and I thank them all. The main contributors over 1996–2006 were Gary McElrea, Lisa McElrea, John Williamson, Nyree Fea, Samantha Brown, Rachel Paterson, Chris Brausch, Louise Chilvers, Phil Knightbridge, Kathryn Knightbridge, Joelle Taillon, Tom Whiteford, Richard Heyward, Peter Berben, and Kev Drew. Nick Spencer maintained the database.

Computations used the New Zealand eScience Infrastructure (NeSI) high performance computing facilities https://www.nesi.org.nz/.
